# Progressive alterations in polysomal architecture and activation of ribosome stalling relief factors in a mouse model of Huntington’s disease

**DOI:** 10.1101/2021.12.13.472357

**Authors:** Eva Martin-Solana, Irene Diaz-Lopez, Yamina Mohamedi, Ivan Ventoso, Jose-Jesus Fernandez, Maria Rosario Fernandez-Fernandez

**Author notes:** Department of Psychiatry, University of Pittsburgh, Pittsburgh, PA, 15213, USA. **Individual email addresses:** E Martin-Solana I Diaz-Lopez Y Mohamedi I Ventoso JJ Fernandez MR Fernandez-Fernandez.

## Abstract

Given their highly polarized morphology and functional singularity, neurons require precise spatial and temporal control of protein synthesis. Alterations in protein translation have been implicated in the development and progression of a wide range of neurological and neurodegenerative disorders, including Huntington’s disease (HD). In this study we examined the architecture of polysomes in their native brain context in striatal tissue from the zQ175 knock-in mouse model of HD. We performed 3D electron tomography of high-pressure frozen and freeze-substituted striatal tissue from HD models and corresponding controls at different ages. Electron tomography results revealed progressive remodelling towards a more compacted polysomal architecture in the mouse model, an effect that coincided with the emergence and progression of HD related symptoms. The aberrant polysomal architecture is compatible with ribosome stalling phenomena. In fact, we also detected in the zQ175 model an increase in the striatal expression of the stalling relief factor *EIF5A2* and an increase in the accumulation of eIF5A1, eIF5A2 and hypusinated eIF5A1, the active form of eIF5A1. Polysomal sedimentation gradients showed differences in the relative accumulation of 40S ribosomal subunits and in polysomal distribution in striatal samples of the zQ175 model. These findings indicate that changes in the architecture of the protein synthesis machinery may underlie translational alterations associated with HD, opening new avenues for understanding the progression of the disease.

## Introduction

Huntington’s disease (HD) is a genetically inherited dominant neurodegenerative disorder caused by expansion of a CAG repeat sequence in the coding sequence of huntingtin protein (HTT) [1]. Symptoms manifest at middle age (35-45 years) and life expectancy after disease onset is usually 15–20 years. Initially, the disease primarily affects the corpus striatum, and neurodegeneration selectively affects medium-sized spiny neurons (MSSNs), which are GABAergic projection neurons [2, 3]. HD patients develop progressive motor dysfunction, cognitive decline, and psychological alterations [4, 5]. Although different pharmacological approaches are being explored, there is no effective treatment for the disease [6, 7]. Recently, promising clinical trials of therapies based on antisense oligonucleotides were halted in phase III, as the potential benefits did not outweigh the risks [8].

The CAG repeat is polymorphic in normal chromosomes and is expanded in HD-mutated chromosomes [9]. The number of repeats influences the age of onset and disease severity. The repeat is located in exon 1 and encodes a polyglutamine sequence (polyQ) that resides in the N-terminal region of the protein. The polyQ sequence in HTT is followed by a proline rich domain (PRD) that is also polymorphic in the human population [10]. HTT is a large 348-kDa protein with a considerable degree of conservation from flies to mammals [10]. However, exon 1 is less evolutionarily conserved than other exons. Interestingly, the PRD is found only in mammals, suggesting recent evolution of the HTT protein [11]. In yeast systems the presence of the PRD reduces the toxicity of exon 1 constructs containing polyQ sequences in the pathological range [12], and also reduces the rate of polyQ aggregation *in vitro* [13–16]. Normal HTT function and the basis for the toxic effects of mutant HTT (mHTT) in the context of HD are still not fully understood [17].

As a consequence of their polarized organization and the particularities of synaptic activity, neurons require precise spatial and temporal control of protein translation, and are therefore particularly susceptible to alterations in the mechanisms controlling protein translation [18]. Dysregulation of cellular mechanisms involved in protein synthesis has been widely implicated in the development and progression of diverse neurological and neurodegenerative diseases [18–21]. Specifically, several studies have shown that mutations in proteins involved in resolving ribosome stalling induce motor dysfunction, neurodegeneration [22], and neuronal cell death in mice [23]. A growing body of evidence points to alterations in mHTT translation and in overall protein synthesis in the context of HD. Surprisingly, mRNAs containing expanded CAG repeats are translated more efficiently than those in the non-pathological range [24]. A higher number of CAG repeats favours the binding of a translation regulation complex containing MID1-PP2A. Consequently, the accumulation of mHTT protein is considerably greater than that of non-pathological HTT. These results reveal an interesting gain of function of the mutant protein at the mRNA level [24]. Several research groups have evaluated translation in the context of HD using the SUnSET method, with conflicting findings. The method is based on the ability of puromycin to label newly synthesized peptides when administrated at low doses. On the one hand, Creus-Muncunill *et al*. reported an increase in puromycin incorporation into striatal cells in brain slices from the R6/1 HD mouse model [25], suggesting that protein translation is increased and that an overall increase in protein translation could constitute a novel pathogenic mechanism in HD. Interestingly, proteomic characterization revealed that the increase in translation specifically affected a set of proteins involved in ribosomal and oxidative phosphorylation, while other proteins implicated in neuronal structure and function were downregulated. On the other hand, Joag *et al.* reported reduced incorporation of puromycin into *Drosophila* cells overexpressing a fragment of HTT containing a 138 polyQ expansion [26]. These results were also reproduced in SYS5 and Neuro2a cells and suggest that expression of a pathogenic mHTT fragment induces a deficit in protein synthesis. Similarly, a recent study reported reduced incorporation of puromycin in heterozygous and homozygous ST*Hdh* cell lines generated from a knock-in mouse model containing 111 CAGs as compared with wild type (wt) counterparts (7 CAGs), suggesting reduced protein synthesis in HD [27]. HTT was proposed to promote ribosome stalling through binding to ribosomes, an effect that is exacerbated by mHTT [27]. Interestingly, yeast cells expressing fragments of pathogenic exon 1 show downregulation in the expression of genes implicated in ribosome biogenesis and rRNA processing and metabolism [28], and hence general repression of protein synthesis. The discrepant findings described above may well be due to the different HD models used, or may be representative of distinct disease scenarios. Overall, these results point to dysregulation of protein synthesis in the presence of mHTT.

Electron tomography allows the study of the 3D subcellular architecture and the organization of molecules in their native cellular or tissue environment with nanometric resolution [29, 30]. It is based on the acquisition of images from a sample at different tilt angles that are subsequently processed and combined to yield a 3D reconstruction or tomogram [31]. Multiple studies have employed electron tomography to investigate the spatial distribution and interaction of ribosomes in their native cellular context in bacteria [32], isolated glioblastoma cells [33], and HeLa cells [34], and more recently as a proof of principle in tissue in *Caenorhabditis elegans* [35]. Rapid-freezing techniques are used to ensure optimal structural preservation of cells and tissues for electron tomography. Specifically, high-pressure freezing (HPF) is used to fully vitrify bulk specimens (up to 200–400 μm thick) through synchronized pressurization and cooling of the sample within milliseconds. Tissue samples can either be processed for observation under cryogenic conditions [35] or freeze-substituted to replace the frozen cellular water with organic solvents and embedded in resins at low temperature to proceed with the analysis at room temperature [30,36]. While fully cryogenic conditions are ideal to preserve the structure in closest-to-native conditions and to achieve the maximum structural resolution, these protocols are not yet used routinely for tissues [35]. Thus, HPF and FS is currently the combination of choice for systematic comparative analysis of ultrastructural tissue alterations in pathological conditions [36].

In the present study, we explored the architecture of neuronal polysomes in their native brain context using 3D electron tomography of HPF/FS striatal tissue obtained from the zQ175 knock-in HD mouse model [37]. We observed progressive remodelling of polysomal architecture resulting in a densely packed conformation compatible with ribosome stalling phenomena. Supporting this hypothesis, we observed increased expression of the stalling release factor *EIF5A2* and an increase in the accumulation of eIF5A1, eIF5A2 and hypusinated eIF5A1 in HD model striatal samples. Polysomal sedimentation gradients revealed a remarkable increase in the accumulation of free 40S ribosomal subunits in striatal samples from HD mice and differences in the distribution of polysomes. These changes in the architecture of the protein synthesis machinery may constitute the basis of translational alterations associated with HD.

## Materials and Methods

### Animals

A stable colony of the zQ175 mice [37] was established using founders donated by the Cure Huntington’s Disease Initiative (CHDI) and obtained from Jackson Laboratory Inc. The zQ175 line is a knock-in line bred on a C57BL/6J background with an endogenous murine *HTT* gene containing a chimeric human/mouse exon 1 with approximately 190 CAG repeats (B6.12951-Htt<tm1Mfc<190JChdi). Heterozygous mice and controls were bred as a stable colony in the animal facility of the Centro Nacional de Biotecnología, Madrid, and provided with food and water *ad libitum*. All experiments complied with Spanish and European legislation and Spanish National Research Council (CSIC) ethics committee on animal experimentation.

### Sample preparation based on HPF/FS for electron microscopy

Brain tissue samples were prepared for electron microscopy (EM) and tomography following our protocols for optimal structural preservation based on high-pressure freezing and freeze-substitution (HPF/FS), as previously described [36]. In short, mouse brains were dissected immediately post-mortem and 200-μm-thick sagittal slices were cut using a tissue slicer (Stoelting, Co.). Striatal samples were promptly extracted, placed on a flat specimen carrier, and frozen under high-pressure in a Leica EMPACT2 device. The samples were further processed with freeze-substitution of frozen water to methanol and embedded in Lowicryl resin HM20 with a Leica AFS2 EM FSP system. Sections (250 nm) were obtained from the resin-embedded samples using a Leica Ultracut EM-UC6 ultramicrotome, and placed on Quantifoil S7/2 grids.

### Electron microscopy

A conventional JEOL JEM-1011 electron microscope (100 kV) was used to screen the 250-nm-thick sections, check the integrity of the tissue samples, and select areas of interest. An average of 5 EM grids per age/genotype were observed. Control and zQ175 animals of 2, 8, 10, and 11 months of age were analysed (2 months: 2 wt males, 1 heterozygous female; 8 months: 1 wt male, 1 heterozygous male; 10 months: 2 wt males, 1 heterozygous male; 11 months: 2 wt males, 2 heterozygous males). Cells compatible with striatal medium-sized spiny neurons were selected based on morphological criteria [38] (Fig. S1) and representative cytoplasmic areas with abundant ribosomes were selected for subsequent 3D studies with ET.

### Electron tomography

Tomographic data were acquired by taking series of images from the sections while tilting them within a range of ±60° at 1° intervals around a single tilt-axis. The tilt-series were acquired using a Thermo Fisher Scientific Talos Arctica electron microscope (200 kV) equipped with a Falcon II electron direct detector or using a FEI Tecnai G2 (200 kV) equipped with a CCD camera. The pixel size at the specimen level was 0.37 nm and 0.59 nm, respectively. For processing, visualization and analysis, images were rescaled with a binning factor of 4. Prior to ET, grids were incubated in a solution of 10-nm diameter colloidal gold (EM.BSA 10, Electron Microscopy Sciences, Hatfield, PA, USA) to facilitate subsequent image alignment. An average of 10 cells were analysed per animal and an average of 16 tomograms taken per animal.

Tilt-series alignment and calculation of the 3D tomograms were conducted using IMOD standard software [39], applying standard protocols [31]. Images of the tilt-series were mutually aligned using the colloidal gold beads as fiducial markers. Tomograms were reconstructed with the standard method, weighted back-projection (WBP), using a filter that simulates an iterative reconstruction method (SIRT) [31].

### Computational analysis of the ribosomal pattern in tomograms

To characterize the ribosomal pattern observed in the tomograms, we developed computational procedures to (1) automatically identify ribosomes and (2) examine how they are organized. The procedure for ribosome identification is based on the Laplacian of Gaussian (LoG), a strategy commonly used in the field of computer vision to detect ‘blobs’. Here, we extended this strategy to 3D in order to work with tomograms. Essentially, the procedure applies Gaussian filtering, with a standard deviation related to the size of the target 3D blob (i.e., the ribosome, around 25 nm in diameter) and, afterwards, a Laplacian operator (i.e., second-order derivative) is applied to the Gaussian-filtered tomogram. The peaks in the resulting LoG map correspond to positions at which ribosomes are located in the tomogram. The final ribosome positions are extracted using a segmentation operation based on thresholding of the LoG map.

To analyse ribosomal organization based on the detected ribosome positions, we implemented second-order spatial point pattern analysis techniques extended to 3D. Specifically, we used Ripley’s K function [40], *K(r)*, which allows us to measure the expected number of ribosomes within a distance *r* from an arbitrary ribosome, normalized by the density of the ribosome distribution. This function can be mathematically expressed as follows:

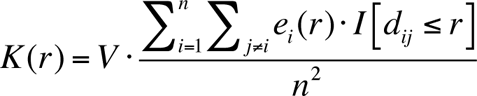

where *V* is the volume of the tomogram, *n* denotes the number of ribosomes in the tomogram, *d_ij_* represents the Euclidean distance from the ribosome *i* to *j*, *I[.]* is an indicator function that equals 1 if the distance *d_ij_* is within the radius *r,* and otherwise equals 0. The term *e_i_(r)* is an edge-correction factor to properly weight the contribution of ribosomes with partial neighbourhood due to their proximity to the edge of the tomogram [40]. Ripley’s K function is calculated at a range of distances *r*, with a recommended maximum value that should be lower than half of the shortest dimension of the sample domain [40, 41]. In this work, the spatial analysis was applied at distances ranging from 25 nm (i.e. ribosome diameter) to 100 nm.

Under the hypothesis of complete spatial randomness (CSR), the K function is equal to the volume of a sphere of radius *r*: *K_CSR_(r) = 4/3×p×r^3^*. The ratio *K(r)/K_CSR_(r)* thus allows analysis of the ribosomal distribution pattern and its classification into the three main categories [42]: aggregation or clustering (if *K(r)/K_CSR_(r) > 1*); regularity or dispersion (if *K(r*)/*K_CSR_(r) < 1*); or random (if *K(r*)/*K_CSR_(r) = 1*). Fig. S2 illustrates the calculation of Ripley’s K function and the three main spatial distribution patterns.

Analysis of the ribosomal distribution was applied to selected tomograms from 10 and 11-month-old animals, acquired on a FEI Tecnai G2 microscope. The larger field of view of this microscope allowed acquisition of regions with an area up to 2.4 µm x 2.4 µm.

### qRT-PCR

Striatal samples from 12-month-old mouse brains were used for gene expression analysis (8 heterozygous zQ175 animals: 4 females, 4 males and 8 corresponding matched wt controls: 4 females, 4 males). The following genes were analysed: *EIF5A1*, *EIF5A2*, *DHPS,* and *DOHH.* The primers used are described in (Table 1).

**Table 1.**
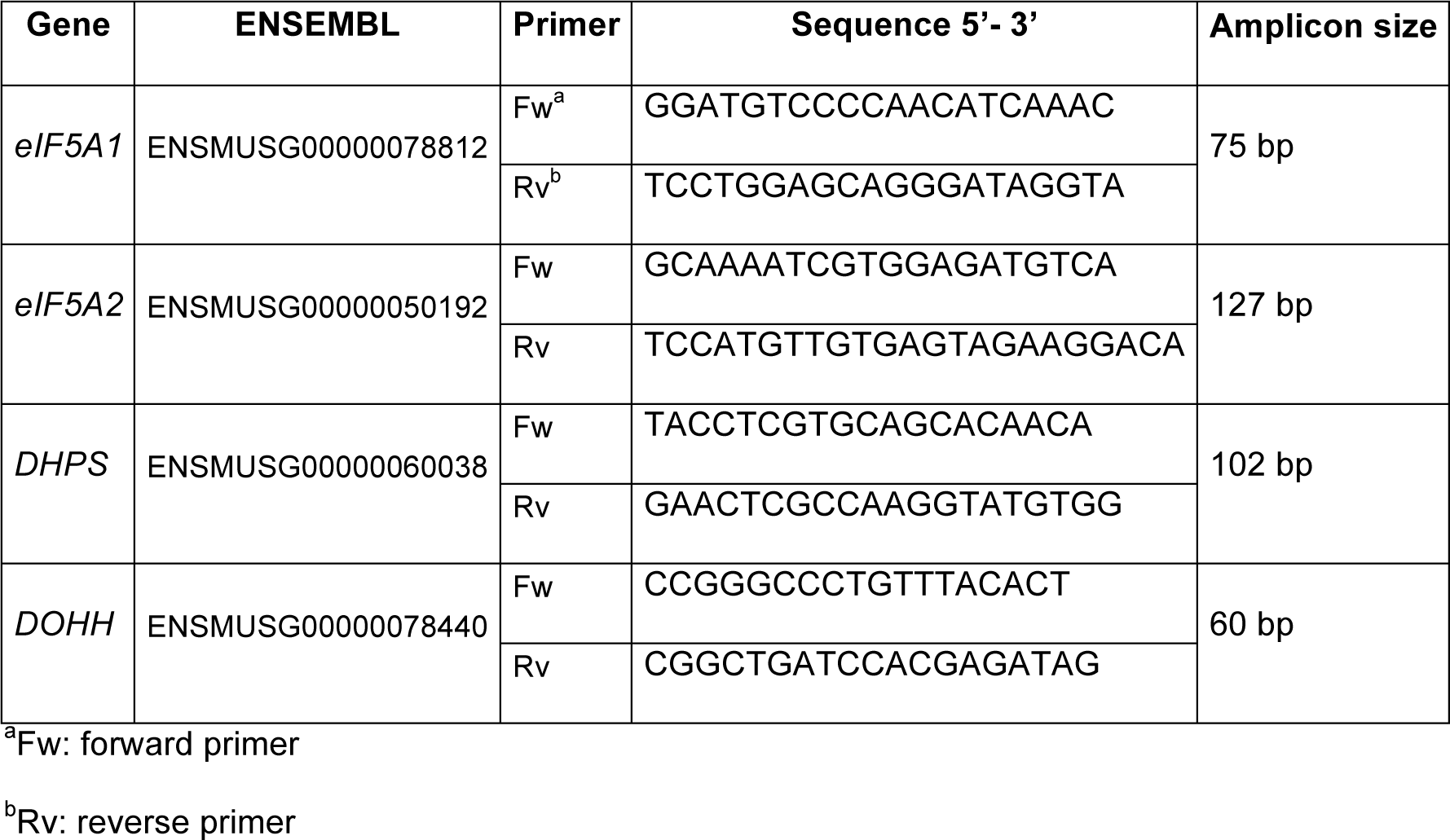
Primers used for qRT-PCR of target genes of interest.

Reference genes were used for normalization of gene expression data (*GAPDH*, *BACT*, *B2M*, *18S,* and *YWHAZ* (Table 2).

**Table 2.**
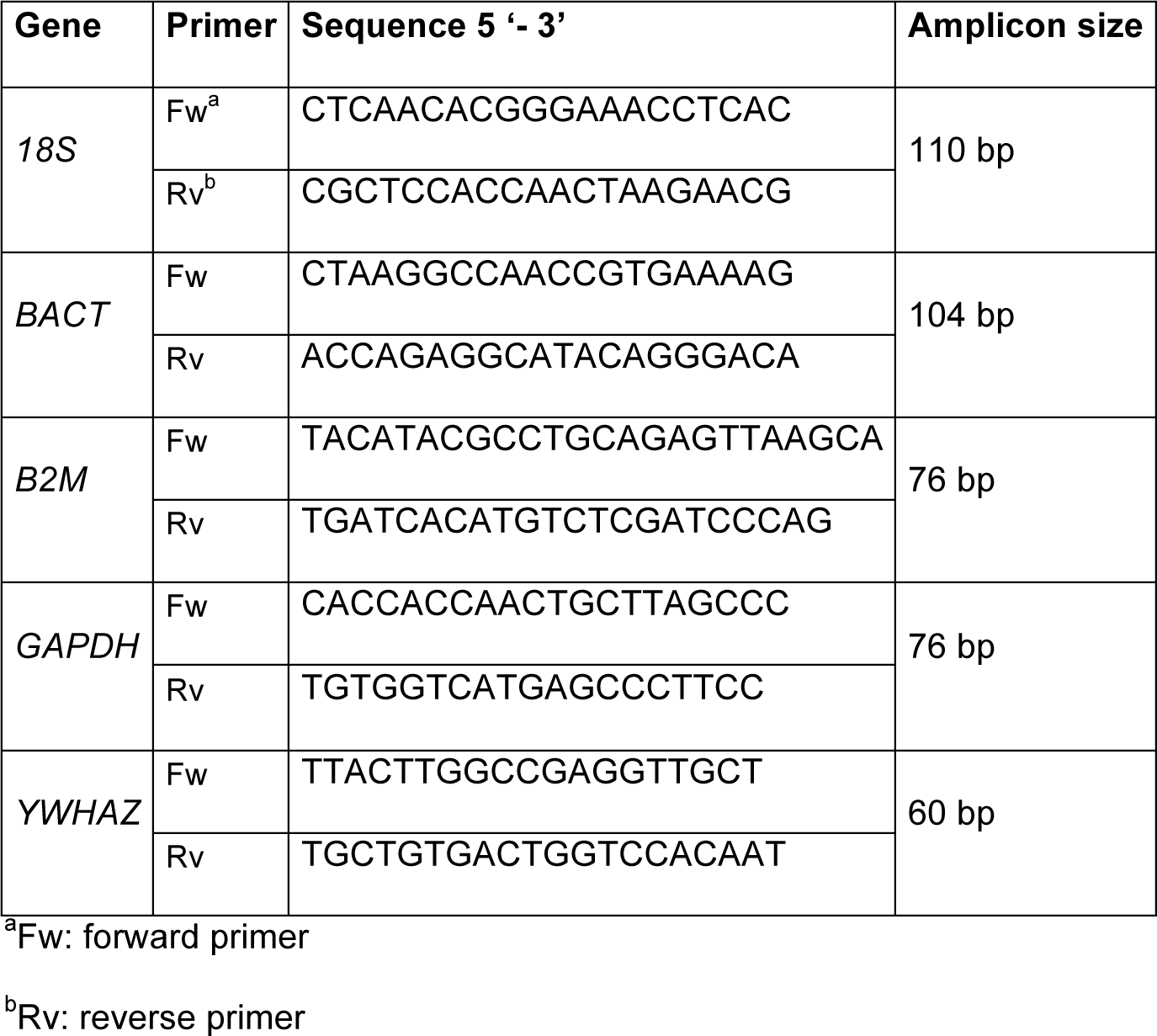
Primers used for amplification by qRT-PCR of reference genes for normalization.

The suitability of reference genes for normalization of gene expression was evaluated using Normfinder algorithm, which showed that *YWHAZ w*as the most stable and accurate gene for normalization. Total RNA was isolated using the Maxwell 16 LEV simply RNA tissue kit (Promega, #AS1280), and RNA integrity was assessed using the Agilent 2100 Bioanalyzer. RIN values ranged from 7 to 9.9, indicating a very high level of sample integrity. cDNA synthesis was performed using the iScript cDNA Synthesis kit (Bio-Rad, #170-8891). qRT-PCR was performed in triplicate on a CFX384 Real Time System C1000 Thermal Cycler (Bio-rad) using Sso Fast EvaGreen Supermix (Bio-Rad, #172-5204). Reactions included a non-template control and primer efficiency curves were generated. ValidPrime kit (TATAA Biocenter, #A105S10) was used as control for genomic background (gDNA). The contribution of gDNA to qPCR signal was less than 6.5% in all cases. A melting curve from 60°C to 95°C (0.5°C/seg) was included at the end of the program to verify the specificity of the PCRs. Data processing was carried out using GenEx v5.4.4 (MultiD Analysis AB, Gothenburg, Sweden). The 2^−Δ(ΔCq)^ method was used to calculate the relative fold change in gene expression of samples normalized to *YWHAZ* gene expression [43].

### Cell culture and transfection

HEK293T cells were cultured in DMEM medium (Sigma, #D6429) supplemented with 10% inactivated foetal bovine serum, non-essential amino acids (1:100, Sigma, #M7145), 0.5 μg/ml amphotericin B (Fungizone^TM^) (Gibco™, #15290018), penicillin-streptomycin (1:100, Sigma, #P4333), and 50 μg/ml gentamycin (Sigma, #G1397). Cells were cultured at 37°C in a 5% CO_2_ atmosphere.

pEGFP-Q23 and pEGFP-Q74 plasmids were a gift from David Rubinsztein (Addgene plasmid # 40261 and 40262) [44]. These express a GFP protein fused to the sequence encoded by part of human *HTT* exon1. Fusion proteins contain either a non-pathological polyQ repeat (23Q) or a polyQ repeat in the pathological range (74Q). These plasmids were co-transfected with pCMV-NP (a plasmid expressing the influenza virus nucleoprotein) as a transfection control [45].

The day before transfection 6×10^5^ cells were plated per well in a M6 plate (Falcon, # 353046) to obtain 70-80% confluence. For actual transfection, plasmids were diluted in a solution containing 250 mM CaCl_2_ to achieve a final amount of 1 or 3 μg per well (final volume per well: 3 ml) for *HTT* plasmids and 2 μg for pCMV-NP. The DNA/CaCl_2_ solution was mixed for 1 min with an equal volume of filtered HBS buffer (50 mM HEPES pH 7.05, 1.5 mM Na_2_HPO_4_, 140 mM NaCl). Cells were harvested in the culture medium after the corresponding period of incubation and centrifuged at 210 *g* for 5 min at 4°C. The pellet was then washed with PBS1x and centrifuged as before. The pellet was frozen and stored at -80°C. Cell lysis was achieved by thermal shock (3 cycles of 15 min at RT+ 30 min at -80°C) in a buffer containing 50 mM Tris-HCl pH 7.4 and 150 mM NaCl. The lysate was centrifuged at 16,100 *g* for 10 min at 4°C and the supernatant used for Western-blot (WB) analysis as described below. Before recovery of the cells, the plates were observed under a Leica DMI6000B microscope at 10x magnification for the detection of GFP-fused proteins.

### Polysome sedimentation gradients

Polysome profiles were generated as previously described [46–48], with some modifications. Briefly, the striatum was dissected from brain and immediately frozen in liquid nitrogen. Samples from mice of the same genotype were pooled to overcome the limitation of the low sample weight (4 mice per genotype: 2 males, 2 females). Frozen samples were ground with a mortar and pestle under liquid nitrogen and homogenized in ice-cold polysome extraction buffer (20 mM HEPES pH 7.4, 150 mM KCl, 5 mM MgCl_2_, 0.5 mM DTT, 1% Triton X-100, 200 µg/ml cycloheximide [CHX], heparin 20 µg/ml, and 40 U/ml RNase inhibitor [New England Biolabs, Cat. no. M0307S]) using a potter homogenizer (10–12 strokes). After three passages through a 23G needle, the homogenates were centrifuged at 2000 *g* for 5 min at 4°C. The supernatants were subsequently centrifuged twice at 16,100 *g* for 10 min at 4°C. The final supernatants (600 μl) were loaded onto 10–50% continuous sucrose density gradients in polysome buffer (20 mM HEPES pH 7.4, 150 mM KCl, 5 mM MgCl_2_) (Gradient Master™, BioComp) and centrifuged in an SW40Ti swing-out rotor (Beckman Coulter) at 250,000 *g* for 2 h at 4°C. Fractions (1 ml) were collected using a density gradient fractionation system (ISCO) with continuous monitoring of absorbance at 254 nm.

Polysomal sedimentation gradients of transfected cells were performed as previously described [49], with some modifications. Briefly, 24 h post transfection, HEK293T cells were treated with 100 µg/ml CHX for 3 min at 37°C. Cells were then washed 3 times with cold CHX (100 µg/ml) in PBS (1x), harvested with residual CHX/PBS and centrifuged at 1000 *g* for 5 min at 4°C. The cell pellet was washed with ice-cold CHX/PBS, centrifuged at 1000 *g* for 5 min at 4°C and resuspended in 500 µl of polysome extraction buffer (100 µg/ml CHX without heparin). The cell lysate was mixed through a 23G needle and incubated on ice for 10 min (vortexing every 2-3 min). The homogenates were centrifuged at 2000 *g* for 5 min at 4°C and subsequently centrifuged twice at 16,100 *g* for 10 min at 4°C. The final supernatants (400 µl) were loaded onto a 10–50% continuous sucrose density gradient (Gradient Station™, BioComp) and centrifuged in an SW40Ti swing-out rotor at 250,000 *g* for 2 h at 4°C. Fractions (1 ml) were collected manually.

### Western-blot analysis

Bulk striatal samples were processed for WB as described previously [50]: seven 15 month old heterozygous animals (2 females, 5 males) and seven corresponding wt controls were used (4 females, 3 males). Samples (25-30 micrograms total protein) were loaded onto 4-20% (Bio-Rad, #4561096) mini-protean TGX precast gels and run in Tris-glycine/SDS running buffer (Bio-Rad, #1610732). Proteins were transferred to nitrocellulose membranes (BioRad, 1620112). For normalization total protein content was quantified by using Revert™ 700 Total Protein Stain (926-11021, LI-COR). Membranes were incubated with the following antibodies: mouse monoclonal raised against eIF5A1 (SAB1402762, Merck), rabbit polyclonal raised against eIF5A2 (17069-1-AP, Proteintech) and rabbit polyclonal anti-hypusine (ABS1064-I, Merck).

Fractions from sucrose gradients were loaded onto NuPAGE 12% Bis-Tris gels (Invitrogen, NP0343) and run in MES running buffer (Invitrogen, NP0002) and transferred as described above. Membranes were incubated with the following primary antibodies: rabbit polyclonal anti-RPS6 (Elabscience, E-AB-32813), anti-RPL7 (Elabscience, E-AB-32805), and anti-hypusine (Creative Biolabs, PABL-202). For accurate quantitative comparisons membranes were first incubated with anti-RPS6 antibody. After completing the WB, membranes were consecutively incubated with anti-RPL7 antibody. For this reason, the residual signal of anti-RPS6 antibody is visible in images of the RPL7 signal.

Samples from transfected HEK293T cells were processed as shown above and membranes were incubated consecutively with anti-GFP (Genetex #GTX113617) and anti-NP [45].

The secondary antibodies used were either IRDye 680RD (925-68071, Li-Cor) and IRDye 800CW (925-32210, Li-Cor) (for bulk striatal samples analysis and eIF5A1/eIF5A2 antibody characterization) or AffiniPure Goat anti-rabbit IgG (H+L) (Jackson ImmunoResearch, 111-035-003) for the rest of WBs. IRDye antibodies were used at 1:2500 and AffiniPure ones at 1:1000 dilutions. The peroxidase reaction for AffiniPure antibodies was developed with Clarity™ Western ECL substrate (Bio-Rad, 1705061). Imaging was performed using either an Odyssey Fc imaging system (Li-Cor) (quantified using Image StudioTM software) or a Molecular ChemiDoc^TM^ Touch Imaging System (Bio-Rad) (quantified using Image Lab 6.0.1 software).

### Statistical analysis

Statistical analyses were performed using GrapPad Prism 8. Normal distribution of the data was assessed using the Shapiro-Wilk test. A two-tailed unpaired Student t-test was used to compare normal distributed data (after assessing the equality of variances using Levene’s test and/or Welch’s correction). The level of significance was set at p<0.05. All data are presented as mean ± SD.

## Results

### Ribosomal/polysomal organization is altered in medium-sized spiny neurons in the striatum of zQ175 mice

HD involves the selective degeneration of striatal MSSNs. Therefore, we examined the architecture of the protein synthesis machinery in its native cellular context, using electron tomography to identify and characterize alterations in these neurons.

Brain tissue samples from HD animal models (heterozygous zQ175) at different ages (2, 8, 10, and 11 months) and the corresponding controls were prepared with HPF/FS for electron microscopy and tomography, as described in the Materials and Methods. This preparation protocol ensures optimal structural preservation of tissue samples. Conventional bidimensional (2D) electron microscopy of 250 nm-thick sections was used to confirm sample integrity and select areas of interest. Representative cytoplasmic areas were selected from the soma of cells compatible with striatal MSSNs where ribosomal and polysomal crowding was identified and examined by electron tomography to determine 3D volumes.

Fig. S3 and S4 show representative EM images and slices of 3D volumes obtained from a wt control and a zQ175 heterozygous mice at advanced age (11 months). In wt samples ribosomes are abundant, are continuously distributed across the entire cytoplasm, and form polysomal clusters in which individual ribosomes can be clearly identified. By contrast, ribosomal density is reduced in the zQ175 model and the polysomes form sparse clusters of tightly-coupled ribosomes with large areas of empty cytoplasmic space between the clusters. Fig. 1 shows representative tomographic slices from animals at different ages: the distinctive pattern in HD mice becomes more pronounced with age. Together, these results indicate that remodelling of polysomal architecture in HD begins at ages (8 months) when some of the relevant phenotypes of this model (diminished rotarod performance, striatal atrophy) [37] become apparent. Moreover, these alterations in polysomal architecture show a clear age-related progression.

**Figure 1.**
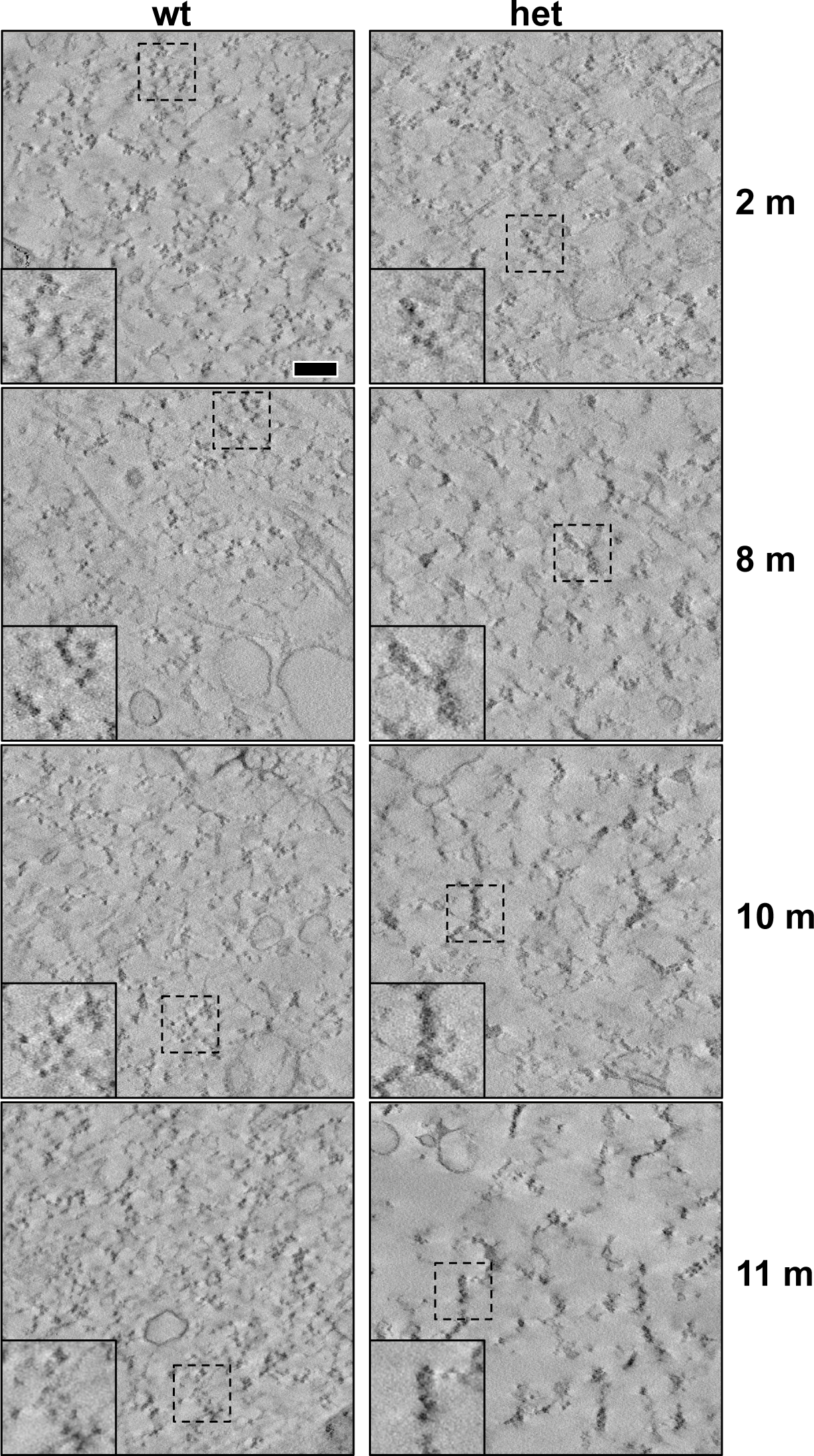
Progressive alterations in ribosomal/polysomal architecture in HD observed with ET. Slices of tomograms from wt (left) and heterozygous (het) zQ175 mice (right). From top to bottom, images correspond to mice aged 2, 8, 10, and 11 months. Insets are magnified views of dashed-line boxes. Scale bar: 200 nm.

To conduct an objective quantitative analysis, we developed a methodology for automated identification of ribosomes in tomograms followed by a second-order spatial analysis using Ripley’s K function, as described in the Materials and Methods section. Fig. S3D and S4D depict the detection of ribosomes in respective tomograms from wt controls and zQ175 mice. These images show that the cytoplasm in wt animals is densely populated with ribosomes whereas in zQ175 mice the ribosomes are organized in groups.

Spatial analysis was applied to the ribosome positions in selected tomograms of different neurons from 10 and 11-month-old animals (Figs. 1 and S5). Ripley’s K-function *K(r)* estimates the number of ribosomes around an arbitrary ribosome within a distance *r*, normalized by ribosomal density. The plots in Fig. 2 depict Ripley’s K-function relative to a random distribution (*K(r)/K_CSR_(r)*, see Materials and Methods). Thus, a unit value indicates a random distribution whereas values >1 imply clustering. The functions show that the ribosomes are significantly more clustered in the heterozygous HD model (*K(r)/K_CSR_(r)* much higher than 1) than in wt controls, especially within shorter distances (up to 50 nm). Additionally, differences in the clustering at 10 and 11-months in the HD model confirm that the phenotype is progressively evolving with age. The wt phenotype also shows some ribosome clustering, though much less pronounced than in the HD model. This is consistent with the fact that ribosomes form polysomes to ensure efficient translation. We also calculated ribosomal density (number of detected ribosomes per volume unit), which was higher in wt controls than the HD model (5163 vs. 2686 ribosomes/µm^3^, analysis combining the 10 and 11-month old animals).

**Figure 2.**
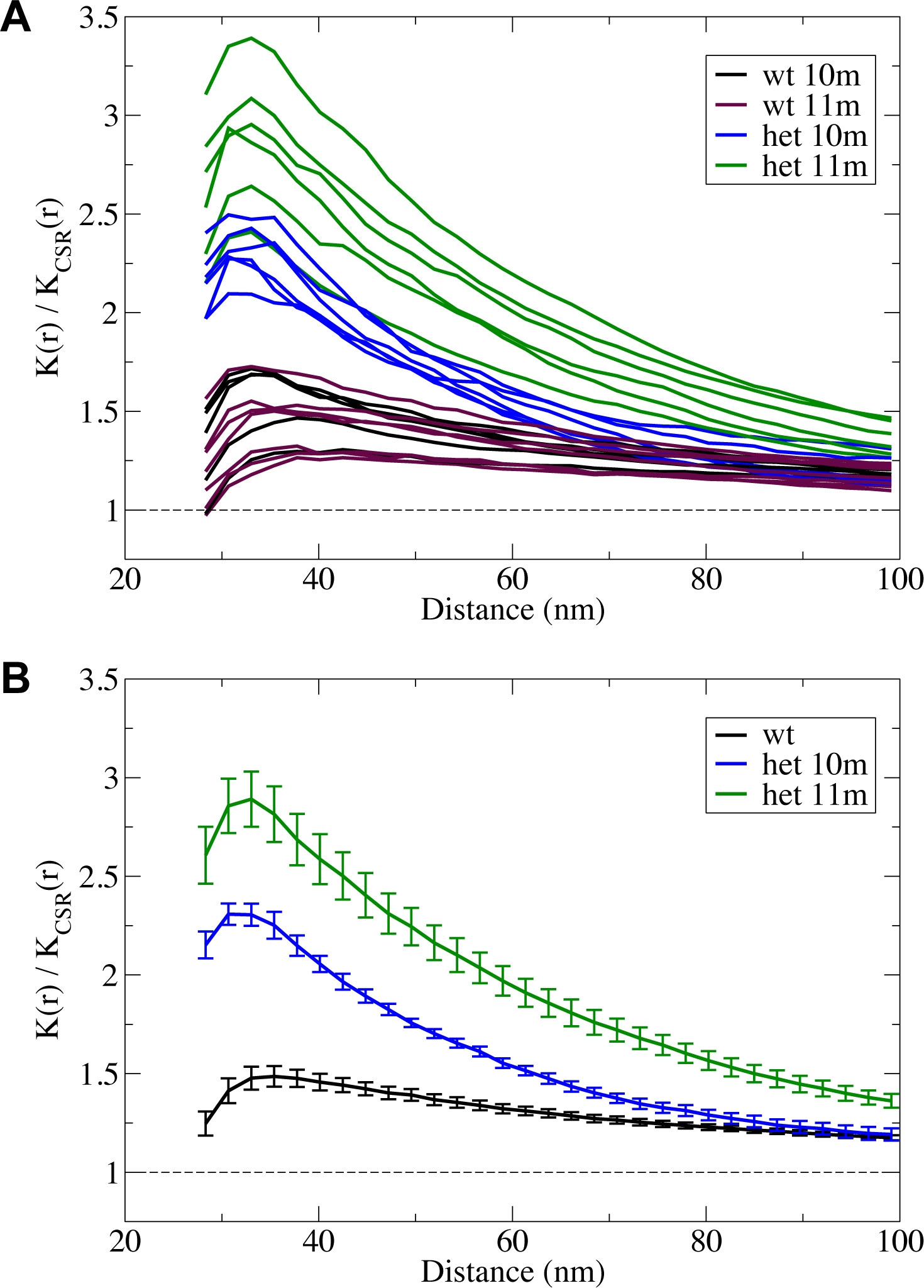
Second-order spatial pattern analysis based on Ripley’s K function. Ripley’s K-function relative to a random distribution, *K(r)/K_CSR_(r)* as a function of the distance calculated for tomograms from different neurons in heterozygous zQ175 and wt mice at ages of 10 and 11 months (A). The plot in (B) depicts mean values and the standard error from the curves of the zQ175 mice at the different ages and also for all wt mice. These Ripley’s K-function plots represent the expected number of ribosomes within a distance r around a random ribosome *(K(r)),* relative to the expected value in a random distribution *(K_CSR_(r*)). Values >1 indicate clustering.

### eIF5A stalling relief factors are activated in the striatum of heterozygous zQ175 mice

The polysomal architecture phenotype of the zQ175 model is reminiscent of ribosome stalling phenomena. It is well established that the presence of consecutive prolines in a polypeptide, as occurs in the PRD region of HTT, induces ribosome stalling. Proline is neither a good donor nor a good acceptor during peptide bond formation, as the reactive amine is embedded in the cyclic lateral chain, thus reducing its nucleophilicity [51–53]. eIF5A and its homologue in bacteria, EF-P, have been shown to participate in stimulating peptide bond formation between consecutive prolines, thereby releasing ribosome stalling [54–56].

eIF5A is encoded by two distinct genes, giving rise to two different homologous proteins: eIF5A1 and eIF5A2 [57]. While eIF5A1 is widely expressed in all tissues, eIF5A2 is weakly expressed in the majority of tissues with the exception of certain areas of the brain and testis, where higher levels are detected [58]. eIF5A proteins are the only known proteins containing the amino acid hypusine, formed by a posttranslational modification of lysine. The role of eIF5A in releasing ribosome stalling is dependent on hypusine [53, 59, 60]. Formation of hypusine involves two enzymatic reactions [60, 61]: deoxyhypusine synthase (DHS) produces deoxyhypusine, which is then hydroxylated by deoxyhypusine hydroxylase (DOHH) to produce hypusine. We investigated whether activation of the eIF5A pathway may be specifically induced in zQ175 mice. Using qRT-PCR, we analysed the expression of eIF5A factors and related genes in the striatum of 12-month old zQ175 mice and corresponding wt controls. As shown in Fig. 3A, we observed no significant difference in the expression of genes encoding eIF5A1, DHPS, or DOHH, and a significant increase in the expression of *EIF5A2* in the striatum of zQ175 heterozygous animals. Overexpression of *EIF5A2* could reflect a response to stalling in HD conditions. It is important to note that the analysis of bulk striatal samples could conceal differences that may be specific to MSSNs.

**Figure 3.**
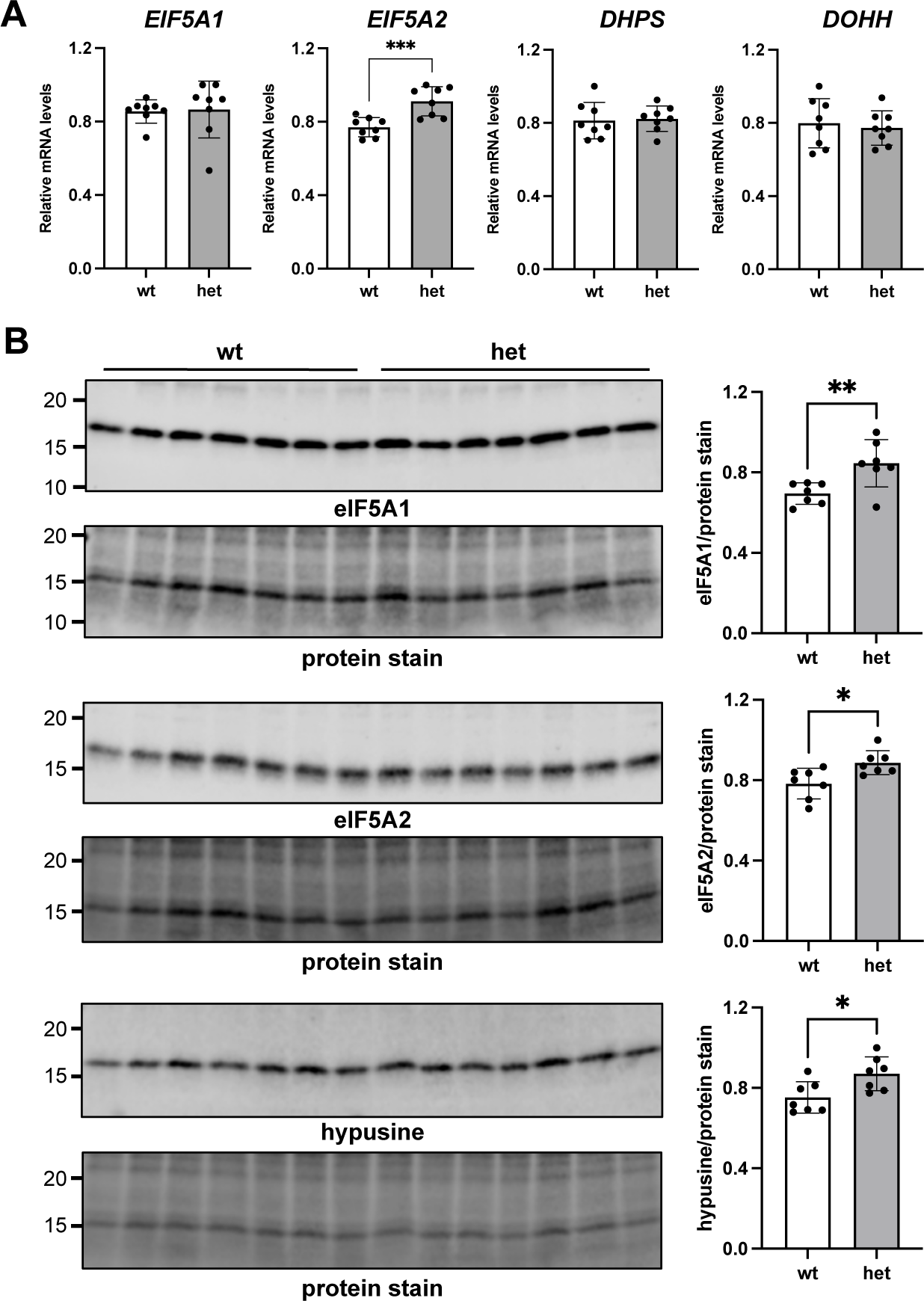
eIF5A stalling relief factors are activated in the striatum of heterozygous zQ175 mice. **(A)** Relative accumulation of *EIF5A1*, *EIF5A2*, *DHPS* and *DOHH* mRNA in the striatum of heterozygous zQ175 mice and wt controls analysed by qRT-PCR. Data are normalized to expression levels of the reference gene *YWHAZ*. For each independent gene the highest value after normalization to *YWHAZ* was set to 1 to facilitate comparisons. Data followed a normal distribution (Shapiro-Wilk test) with the exception of *EIF5A1*. No significant difference between groups was observed for *EIF5A1, DHPS,* or *DOHH* gene expression (p=0.2786, 0.8130, and 0.6642, respectively). *EIF5A2* expression was significantly increased in heterozygous zQ175 mice versus controls (p=0.0009). Data are presented as the mean ±SD (n=8). **(B)** Western-blot analysis of the accumulation of eIF5A1, eIF5A2 and hypusinated eIF5A factor in the striatum of 15-month-old heterozygous zQ175 mice and wt controls. In the eIF5A2 blot the shadow band below the eIF5A2 corresponds to the antibody cross-reaction with eIF5A1 (see Fig. S6). Total protein in each lane was quantified by staining with Revert™ 700 Total Protein Stain (926-11021, LI-COR) for normalization of loading. The panel in the figure shows only the area in the molecular weight range of the proteins of interest. Graphs on the right show the quantification of relative accumulation. In all cases data followed a normal distribution. We detected a significant increase in the accumulation of eIF5A1 (p=0.0095), eIF5A2 (p=0.0149) and hypusinated eIF5A1 (p=0.0181) in the heterozygous animals. Data are presented as the mean ±SD (n=7).

We next examined the accumulation of eIF5A1, eIF5A2 and hypusinated eIF5A forms in striatal samples by WB. eIF5A1 and eIF5A2 have an 84% identity at the amino acid level, consequently many antibodies raised against one of them cross-react with the other. To unequivocally distinguish both isoforms we evaluated the specificity of the antibodies used in this work against recombinant eIF5A1 and eIF5A2 expressed in bacteria (Fig. S6).

We observed a significant increase in the relative accumulation of eIF5A1, eIF5A2 and the hypusine containing form of eIF5A1 in the striatum of heterozygous zQ175 mice versus corresponding wt controls at 15 months of age (Fig. 3B). We were not able to detect in this WB a band corresponding to the hypusine containing form of eIF5A2 (Fig. S6B).

Next, we investigated whether there is any differential recruitment of the active (hypusinated) form of eIF5A1 to ribosomes. We optimized a protocol to prepare polysomal sedimentation gradients: given the low weight of the mouse striatum the protocol was performed using a pool of samples from 4 animals. Fig. 4A shows the profiles obtained for the wt and heterozygous zQ175 pools. The peaks corresponding to 40S (small subunit), 60S (large subunit), 80S (monosome), and polysomes are clearly identifiable in the profiles. Analysis of the fractions by WB (Fig. 4B) confirmed the expected distribution of ribosomal proteins corresponding to small and large subunits (RPS6 and RPL7), respectively. For both wt and heterozygous zQ175 mice, most of the hypusinated eIF5A (>80%) was detected in fractions 1 and 2, which do not contain ribosomal subunits. Thus, most of the active stalling relief factor is not associated to ribosomes. This result could be consistent with a transient interaction of the factor with the 60S subunit [62], or could be indicative of functions of the eIF5A factor other than facilitating elongation in polyproline-induced stalling, as previously proposed [59, 63–65].

**Figure 4.**
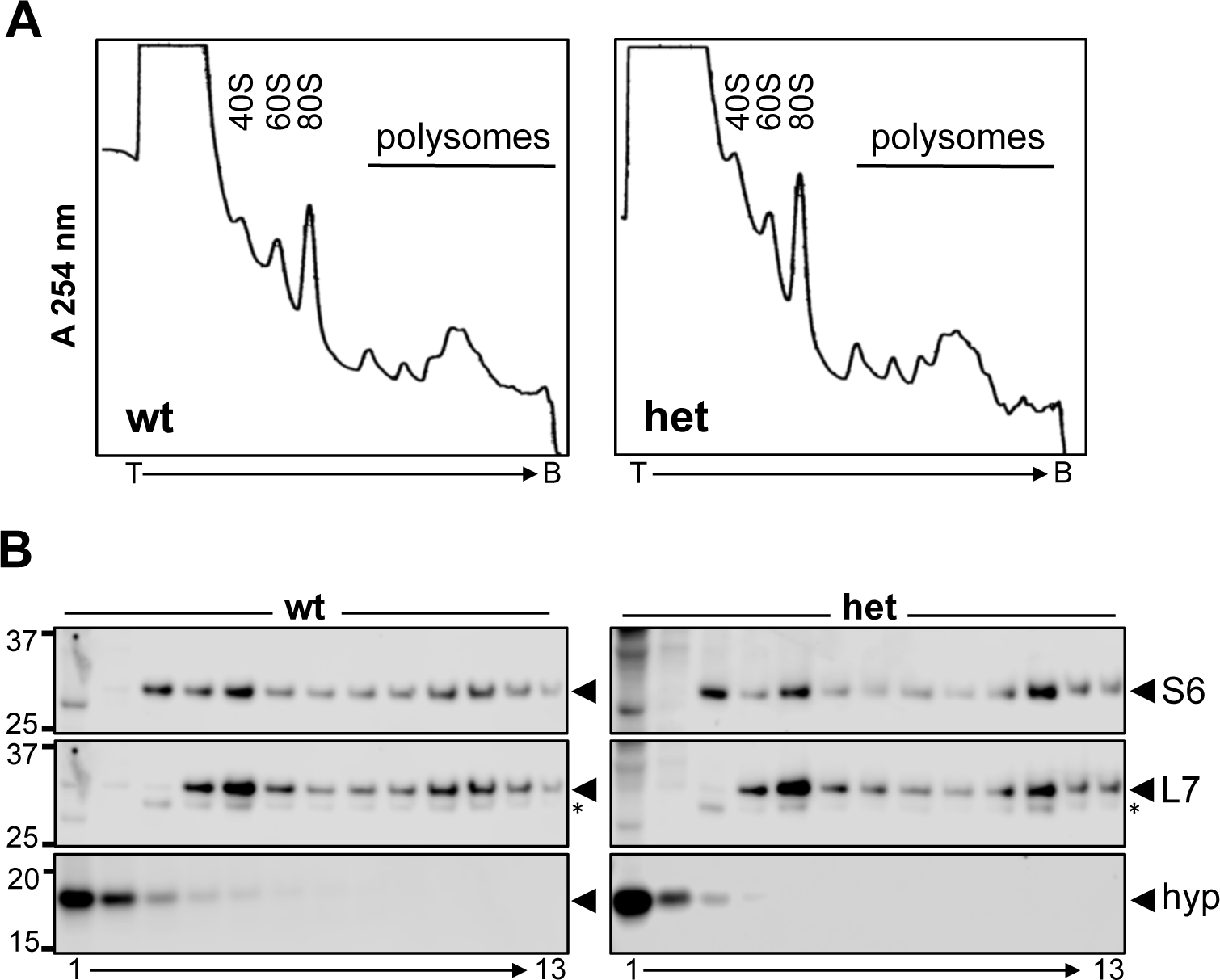
Polysomal sedimentation gradients generated from striatal samples from 9-month-old wt and heterozygous zQ175 mice. **(A)** Profiles obtained by continuous monitoring of absorbance at 254 nm. Peaks corresponding to 40S, 60S, 80S, and polysomes are identifiable. T indicates the top and B the bottom of the gradient. **(B)** Accumulation of RPS6 (S6), RPL7 (L7) and hypusinated eIF5A factor in gradient fractions. Fraction 1 corresponds to the top and fraction 13 to the bottom of the gradient. Most hypusinated eIF5A (>80%) accumulates in the first two fractions that do not contain ribosomal subunits (soluble fractions). Asterisks indicate the residual signal of RPS6.

The profiles of polysomal sedimentation gradients (Fig. 4A) were largely similar for wt and heterozygous zQ175 mice: we observed no remarkable differences in the polysomal area that could correspond to the polysomal compaction phenotype observed by ET. However, the profiles obtained suggest a higher 40S:80S ratio in heterozygous zQ175 versus wt mice. Analysis of fractions by WB (Fig. 4B) showed that the ratio of RPS6 accumulation in fraction 3 (40S) versus fraction 5 (80S) is higher in the heterozygous zQ175 (1.01) than in the wt (0.77), thereby confirming an increased accumulation of free 40S ribosomal subunit in the striatum of heterozygous zQ175 animals. Interestingly, the WB analysis (Fig. 4B) also showed a different distribution of ribosomal proteins in the polysomal area. In the heterozygous zQ175 gradient we observed a maximum peak in fraction 11 (for both RPS6 and RPL7), while in the wt gradient the proteins were more widely distributed across surrounding fractions, with equivalent levels detected in fractions 10 and 11. The relevance of this differential distribution of polysomes and its relation to the architectural phenotype detected by ET remains to be determined.

### The expanded CAG repeat does not have a negative effect on protein accumulation

Next, we explored whether differences in the ribosomal and polysomal organization observed in the striatum of zQ175 mice models are a direct consequence of mHTT expression. We transfected HEK293T cells with plasmids to overexpress GFP protein fused to the sequence encoded by part of the *HTT* exon1 containing either 23 (pEGFP-Q23) or 74 (pEGFP-Q74) CAG repeats, which represent non-pathological and pathological conditions, respectively [44]. Different plasmid concentrations were used (1 μg and 3 μg per well). As a control of transfection, cells were co-transfected with the pNP plasmid (pCMV-NP) [45]. Results in Fig. 5A and 5B show that for both plasmids higher concentrations are associated with greater accumulation of the fusion proteins. Interestingly, at both plasmid concentrations accumulation of the GFP-Q74 fusion protein was greater than that of GFP-Q23 (with a significant effect observed for 3 μg). A similar effect was previously reported [24] in transfection experiments using mouse embryonic fibroblasts and plasmids that express 17Q and 49Q tracks in the context of a wider N-terminal HTT fragment (first 500 amino acids). Those authors observed no differences in the expression of mRNAs encoding Q17 and Q49. They also showed that increasing the number of CAG repeats favours binding of the MID1-PP2A-containing translation regulation complex, thus supporting the view that differences in translation efficiency underlie the protein accumulation effect. Our results confirm that this effect is highly dependent on the CAG sequence coding for the polyQ, as the plasmids we used contain the CAG repeat in the context of a partial and short exon 1 sequence (polyQ is flanked in the N- and C-terminus by 10 and 17 aa encoded by exon 1, respectively).

**Figure 5.**
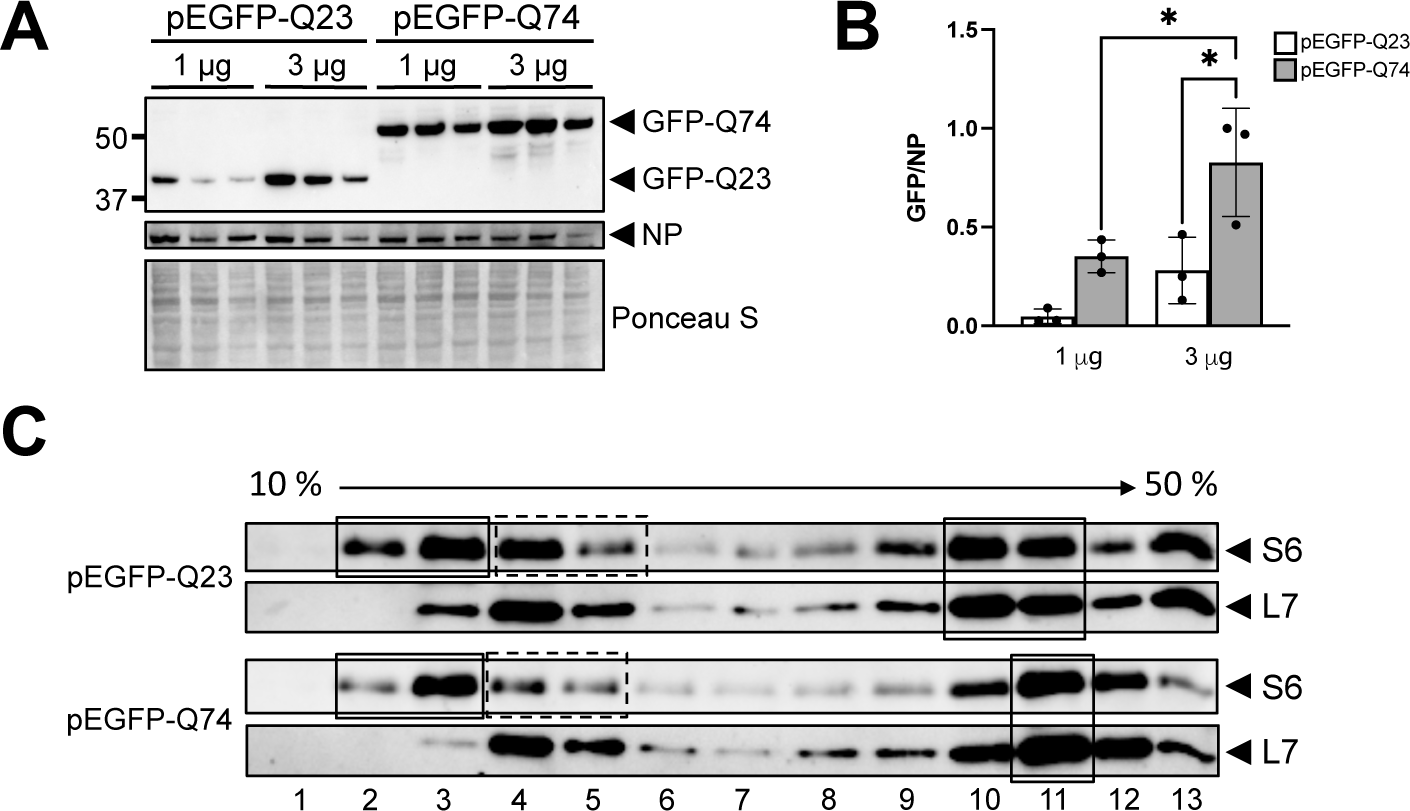
Transfection of HEK293T cells with plasmids expressing a GFP fusion protein containing part of the exon1 *HTT* coding sequence including 23Q or 74Q. **(A)** Accumulation of fusion proteins GFP-Q23 and GFP-Q74 18 hours post-transfection. Middle panel shows NP protein accumulation as a transfection control. The bottom panel shows the Ponceau S staining. Samples were run in triplicate. **(B)** Quantification of the relative accumulation of GFP-Q23 and GFP-Q74 for the different transfection conditions shown in **A**. Mean ± SD values are shown. Increased accumulation of GFP-Q74 was observed at both plasmid concentrations: significant at 3 μg (p=0.0422) but not at 1 μg (p=0.1000). For both plasmids an increase in the amount of transfected plasmid was associated with increased accumulation of the fusion protein (not significant for GFP-Q23 (p=0.1000) but for GFP-Q74 (p=0.0453)). The pEGFP-Q23 1 μg dataset showed a non-normal distribution, all other groups had a normal distribution. **C)** WB analysis of fractions from polysomal sedimentation gradients obtained from HEK293T cells transfected with either pEFGP-Q23 or pEGFP-Q74. Cells were recovered 24 hours post-transfection and the percentage of transfection was 20-25%. Solid boxes on the left highlight the accumulation of RPS6 (small ribosomal subunit) in the fractions of the gradient likely corresponding to 40S subunit and the dashed boxes those corresponding to 80S subunit. The 40S/80S ratio of RPS6 accumulation is 1.4 for GFP-Q23 and 2.8 for GFP Q74, confirming that overexpression of GFP-74Q fusion protein is associated with increased percentage of free 40S subunits. Boxes on the right highlight the differential accumulation of polysomes in the mid polysomal area. Membranes were first incubated with anti-RPS6 and then reincubated with anti-RPL7.

### Overexpression of constructs containing CAG repeats in the pathological range is sufficient to induce increased accumulation of free 40S ribosomal subunits

We next explored whether differences in the polysome sedimentation gradients observed in striatal samples from heterozygous zQ175 mice are a direct effect of mHTT expression. We performed polysomal sedimentation gradient analyses using pEFGFP-Q23- and pEGFP-Q74-transfected cells 24 hours post-transfection. This was done applying the modifications described in the Materials and Methods, and therefore there is a different distribution of the ribosomal species in the fractions of the gradient as compared to gradients performed with striatal samples. As shown in Fig. 5C the differential accumulation of ribosomal proteins, RPS6 and RPL7, indicates that RPS6 accumulation in fractions 2 and 3 is representative of 40S subunit while that in fractions 4 and 5 is representative of 80S. The gradients from cells overexpressing GFP-Q74 (pathogenic range) revealed changes similar to those observed in striatal samples from heterozygous zQ175 mice. Specifically, we could determine that GFP-Q74 overexpression induces increased accumulation of free 40S subunit (Fig. 5C). Additionally, differences observed in the polysomal area mirrored those detected in gradients of striatal samples. While GFP-Q74-overexpressing cells showed maximum accumulation around fraction 11, ribosomal proteins were more widely distributed across surrounding fractions in GFP-Q23-overexpressing cells.

## Discussion

One of the most intriguing characteristics of HD is its slowly progressive nature. While the mutant protein is expressed throughout life, symptoms only begin to manifest at middle age (35-45 years of age), and subsequent life expectancy is 15-20 years. Subtle alterations at the cellular level have been proposed to accumulate overtime, only exerting detrimental effects at later stages [10]. Our electron tomography results indicate progressive disturbance of neuronal polysomal architecture in the striatum of the zQ175 mouse model of HD. In this model, motor coordination phenotypes (measured by rotarod performance) and striatal atrophy become evident from 8 months of age [37]. Interestingly, 8 months was the earliest time point at which rearrangements of polysome structure were evident in our study. This coincidence and the importance of protein synthesis for cellular homeostasis, suggest that the polysomal alterations described here may be relevant to the development of HD-related phenotypes and may reflect disease progression. The protein synthesis disturbances associated with HD [25–27] have led some authors to propose that translational dysfunction contributes to the pathogenesis of HD and that new therapies targeting protein synthesis could help alleviate disease symptoms [26]. Our results further underscore the role of translational dysfunction in HD and point to alterations in the architecture of the protein synthesis machinery as a starting point for translational dysfunction.

Studies using gene-specific deletion approaches have enabled the identification of yeast strains in which mHTT toxicity is suppressed [28]. Interestingly, mHTT overexpression in these strains results in increased expression of the stalling relief factor *EIF5A2*. The increased *EIF5A2* expression that we detected in the striatum of heterozygous zQ175 animals could represent a cellular response to mHTT toxicity. Increased *EIF5A2* expression, increased accumulation of eIF5A1, eIF5A2 and hypusinated eIF5A1 together with progressive compaction of polysomes support a scenario of ribosome stalling induced by mHTT.

eIF5A factors are mainly involved in releasing the ribosome stalling induced by the translation of consecutive prolines [54–56]. Interestingly, the polyQ sequence in HTT is followed by a PRD with abundant consecutive prolines. This PRD is present in both wt and mutant HTT, begging the question why stalling induced by the translation of the PRD is only evident in heterozygous animals. An interesting study reported that the dependence of stalling release on eIF5A factors correlates with the translation initiation rate of each particular mRNA [66]. Thus, when initiation rates are low, translation through polyproline sequences resumes without impeding the progression of upstream ribosomes, even in the absence of eIF5A factors. However, transcripts with a high initiation rate will often engage simultaneously with several ribosomes, which are likely to stall when translating the polyproline sequence and become dependent on the action of eIF5A factors. This is particularly relevant in the context of the findings of Krauß *et al.,* who showed that a higher number of CAG repeats favours binding of the MID1-PP2A-containing translation regulation complex, resulting in more efficient translation [24]. Together, these findings suggest that mHTT mRNA may be more sensitive to the effect of polyproline-mediated stalling because it is more efficiently translated and therefore more dependent on the action of eIF5A relief factors.

We describe excess accumulation of free 40S subunits in the striatum of the heterozygous zQ175 mouse model of HD. The stability and accumulation of small and large ribosomal subunits are coupled as part of a process that may have evolved to ensure adequate translational performance. An excess of 40S subunits could lead to the formation of translation initiation complexes that cannot be converted to fully translating 80S ribosomes and consequently sequester mRNAs, resulting in general disruption of protein synthesis [67].

## Conclusions

Our findings underscore the role of translational dysfunction in HD and point to alterations in the architecture of the protein synthesis machinery in the basis of such dysfunction. Further studies are needed to understand the underlying mechanisms by which mutations associated to HD induce these alterations.

## Abbreviations

2D: Bi-dimensional
3D: Three-dimensional
CCD: Charge-coupled device
CSR: Complete spatial randomness
DHS: Deoxyhypusine synthase
DOHH: Deoxyhypusine hydroxylase
EIF5A: Eukaryotic Translation Initiation Factor 5A
EM: Electron microscopy
ET: Electron tomography
FS: Freeze-substitution
GABA: Gamma-aminobutyric acid
HD: Huntington’s disease
HPF: High-pressure freezing
HTT: Huntingtin protein
LoG: Laplacian of Gaussian
mHTT: Mutant huntingtin protein
MSSNs: Medium-sized spiny neurons
polyQ: polyglutamine
PRD: Proline rich domain
qRT-PCR: Quantitative reverse transcription polymerase chain reaction
RPS6: 40S ribosomal protein S6
RPL7: 60S ribosomal protein L7
SD: Standard deviation
SIRT: Simultaneous iterative reconstruction technique
SUnSET: Surface sensing of translation
WB: Western blot
WBP: Weighted backprojection
wt: Wild type

## Declarations

### Ethical approval and consent to participate

All experiments complied with Spanish and European legislation and Spanish National Research Council (CSIC) ethics committee on animal experimentation.

### Consent for publication

All authors have approved of the contents of this manuscript and provided consent for publication.

### Availability of data and materials

The data of this study are available from the corresponding authors on reasonable request.

### Authors’ contributions

(CRediT: https://onlinelibrary.wiley.com/doi/epdf/10.1002/leap.1210)

EMS: Investigation, Validation, Formal Analysis, Methodology, Writing - Original draft.

IDL: Investigation, Resources, Writing - Review & Editing.

YM: Investigation

IV: Investigation, Resources, Writing - Review & Editing.

JJF: Conceptualization, Methodology, Software, Investigation, Formal Analysis, Writing - Original draft, Funding Acquisition.

MRFF: Conceptualization, Methodology, Investigation, Validation, Formal Analysis, Writing - Original draft, Funding Acquisition.

All authors read and approved the final manuscript.

## Acknowledgements

We thank Pablo Sola Alarcón for valuable technical assistance. We would also like to thank Eber Martínez and all staff at the Animal Facility of the Centro Nacional de Biotecnología (CNB-CSIC, Madrid) for their work over the years. qPCR experiments were conducted at the Genomics and NGS Core Facility at the Centro de Biología Molecular Severo Ochoa, which is part of the CEI UAM+CSIC, Madrid. HPF and FS experiments together with the observation of brain sections were performed in the Electron Microscopy Facility at the CNB-CSIC. The acquisition of tilt series for tomography was performed in the Cryoelectron Microscopy Facility at CNB-CSIC. We are profoundly grateful to the Fundación Ramón Areces for the grant CIVP18A3892 that made this work possible, and to AEI (SAF2017-84565-R, TED2021-132020B-I00, PID2022-139071NB-I00) and Huntington**’**s disease Society of America (HDSA) (Human Biology project 2022) for additional funding. We would also like to thank the Cure Huntington**’**s disease initiative (CHDI) for providing the initial founders of the zQ175 colony within the context of a previous project.

---

**Supplementary Figure S1.**
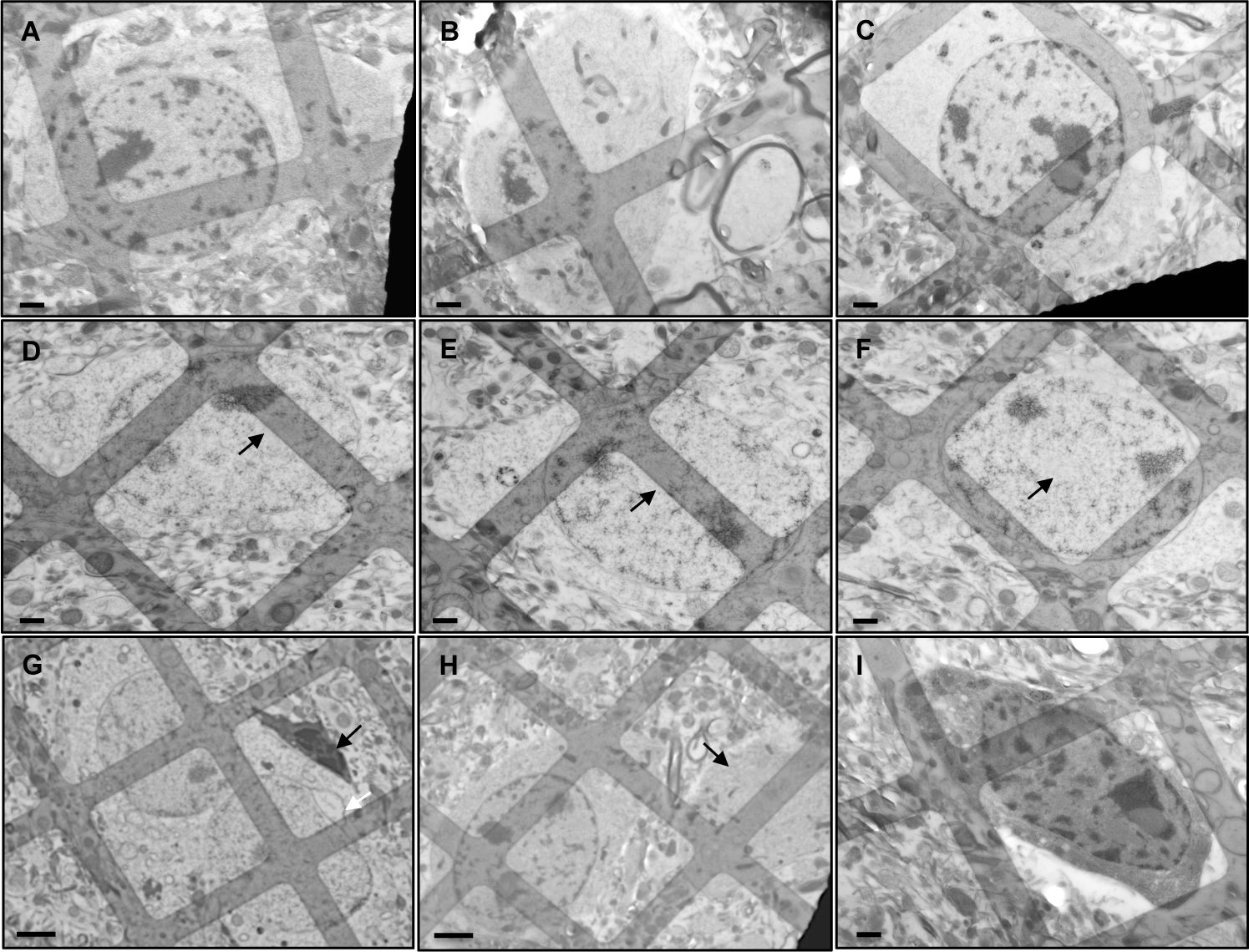
Representative EM images of striatal cells from 250-nm-thick sections of brain tissue. **(A-F)** Cells compatible with medium-sized spiny neurons (MSSN) based on morphological criteria in wt control **(A-C)** and heterozygous zQ175 **(D-F)** mice aged 11 months. In the HD model, these cells appear paler, an effect evident in the chromatin, nucleus and cytoplasm. In addition, nuclear inclusions are identifiable in the HD model (arrows in D-F). **(G)** A cell, likely microglia, with nucleus and cytoplasm that are highly dense (black arrow) is observed near two pale cells compatible with MSSNs. One of these MSSNs contains a nuclear bleb (white arrow), which are often found in HD neurons and are indicative of neurodegeneration. **(H)** A putative MSSN and, on the right, an area of cytoplasm from another cell (arrow) with a similar appearance but not showing the nucleus. **(I)** A cell from a control animal, with size and morphology compatible with a MSSN, with highly dense nucleus and cytoplasm (signs of cell death). Scale bars: 1 μm (A-F, I), 2 μm (G-H).

**Supplementary Figure S2.**
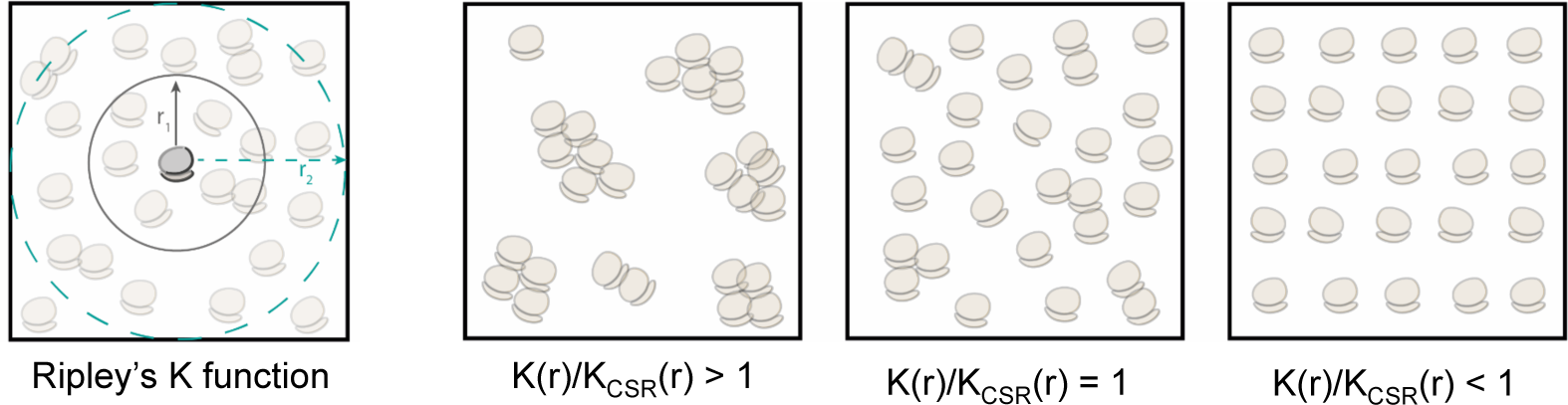
Ripley’s K function and spatial distribution patterns. Left: Ripley’s K function is calculated at a range of distances *r* and measures the expected number of ribosomes within the distance *r* from an arbitrary ribosome, normalized by the density of the ribosome distribution. In this study, the K function has been implemented in 3D, though for simplicity the sketch depicts 2D. Right: Three main spatial distribution patterns can be identified when comparing the actual K function with the expected value under complete spatial randomness (K_CSR_): clustering (K(r)/K_CSR_(r) > 1); randomness (K(r)/K_CSR_(r) = 1); and regularity or dispersion (K(r)/K_CSR_(r) < 1).

**Supplementary Figure S3.**
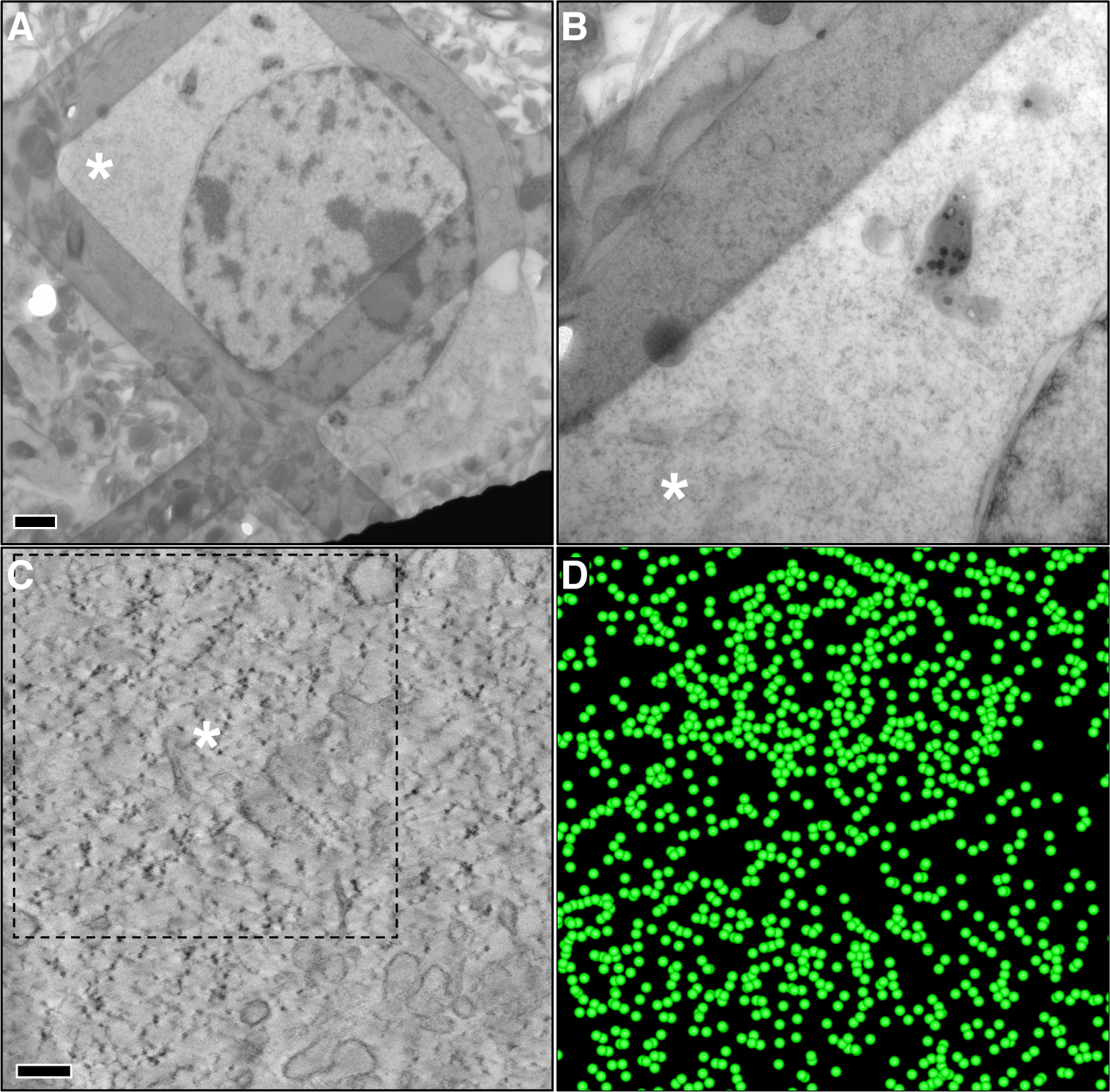
Electron tomography of a striatal neuron from a 250 nm thick section of brain tissue from an 11-month-old wild-type mouse. **(A)** Image from conventional 2D electron microscopy, showing the nucleus of the neuron and cytoplasm. Scale bar: 1 µm. **(B)** Magnified view from the area marked by the asterisk in **A**. **(C)** Tomogram slice computed from a tilt-series acquired at the position marked with the asterisk in **B**. The individual ribosomes are clearly identifiable as black dots in the spotty cytoplasm. The cytoplasm is crowded by numerous ribosomes, and polysomal clusters are also observed. Scale bar: 200 nm. **(D)** 3D visualization of the ribosomes detected using our automated procedure based on Laplacian of Gaussian (LoG) and represented by green spheres. These ribosomes correspond to the area outlined with a dashed line in **C**.

**Supplementary Figure S4.**
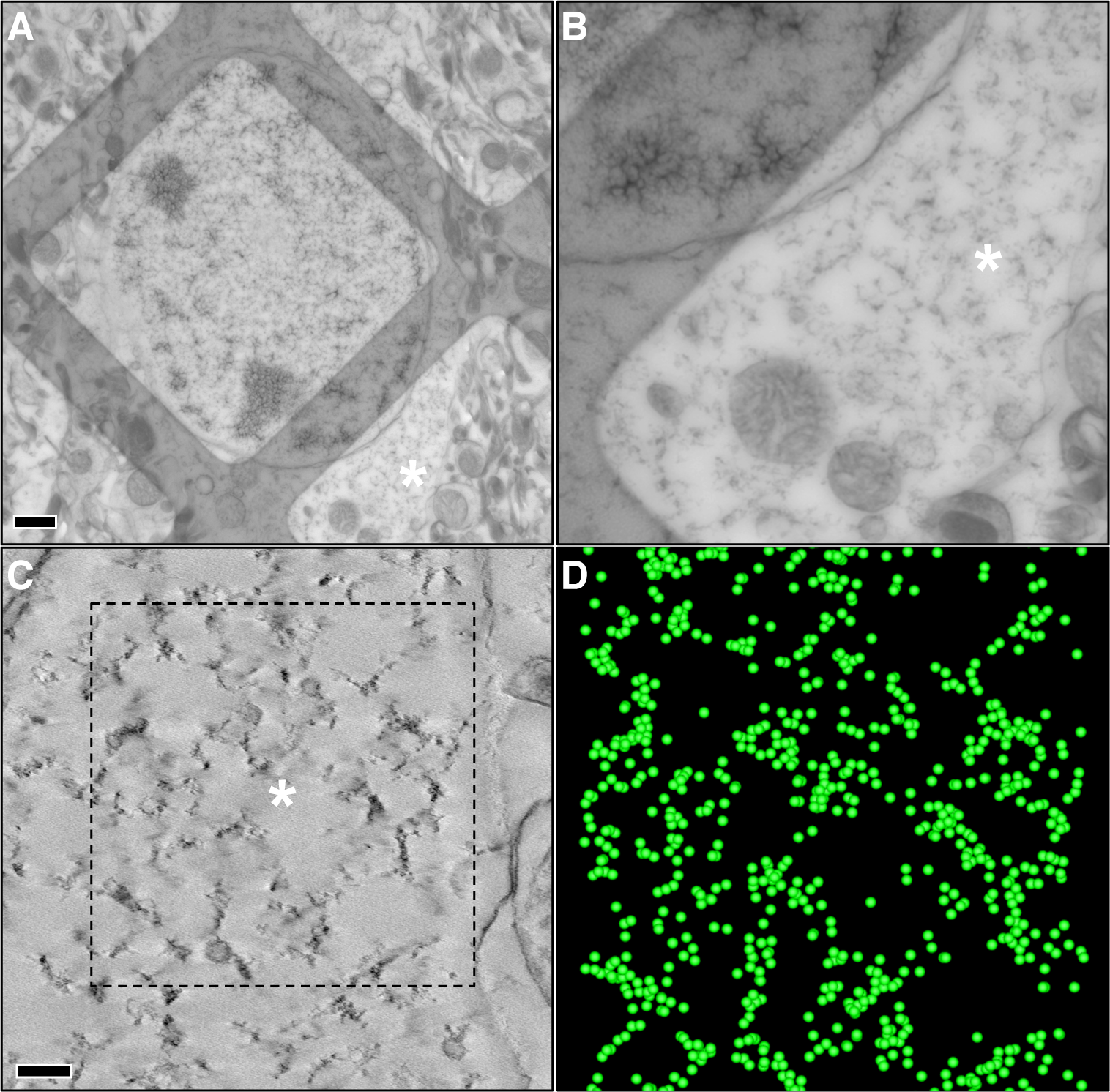
Electron tomography of a striatal neuron from a 250-nm-thick section of brain tissue from an 11-month-old heterozygous zQ175 mouse. **(A-D)** Interpretation of the panels as described in Figure S3. In this case, few individual ribosomes are observed in the tomogram **(C)**. Instead, tightly-coupled clusters of ribosomes are evident, and there is substantial empty cytoplasmic space among the clusters.

**Supplementary Figure S5.**
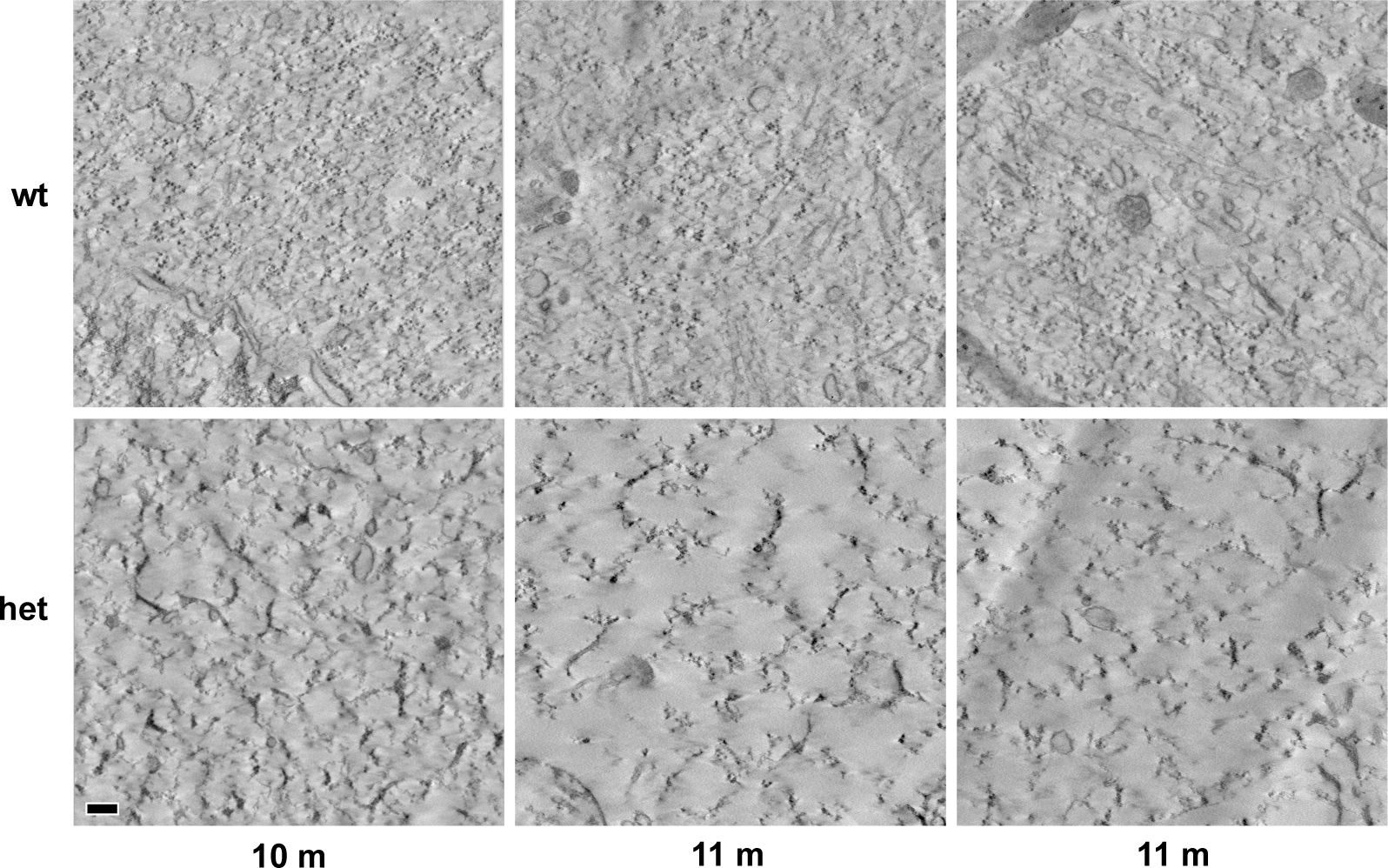
Representative slices of tomograms from 10 and 11-month-old wild-type control (top) and heterozygous zQ175 (bottom) mice. Scale bar: 200 nm.

**Supplementary Figure S6.**
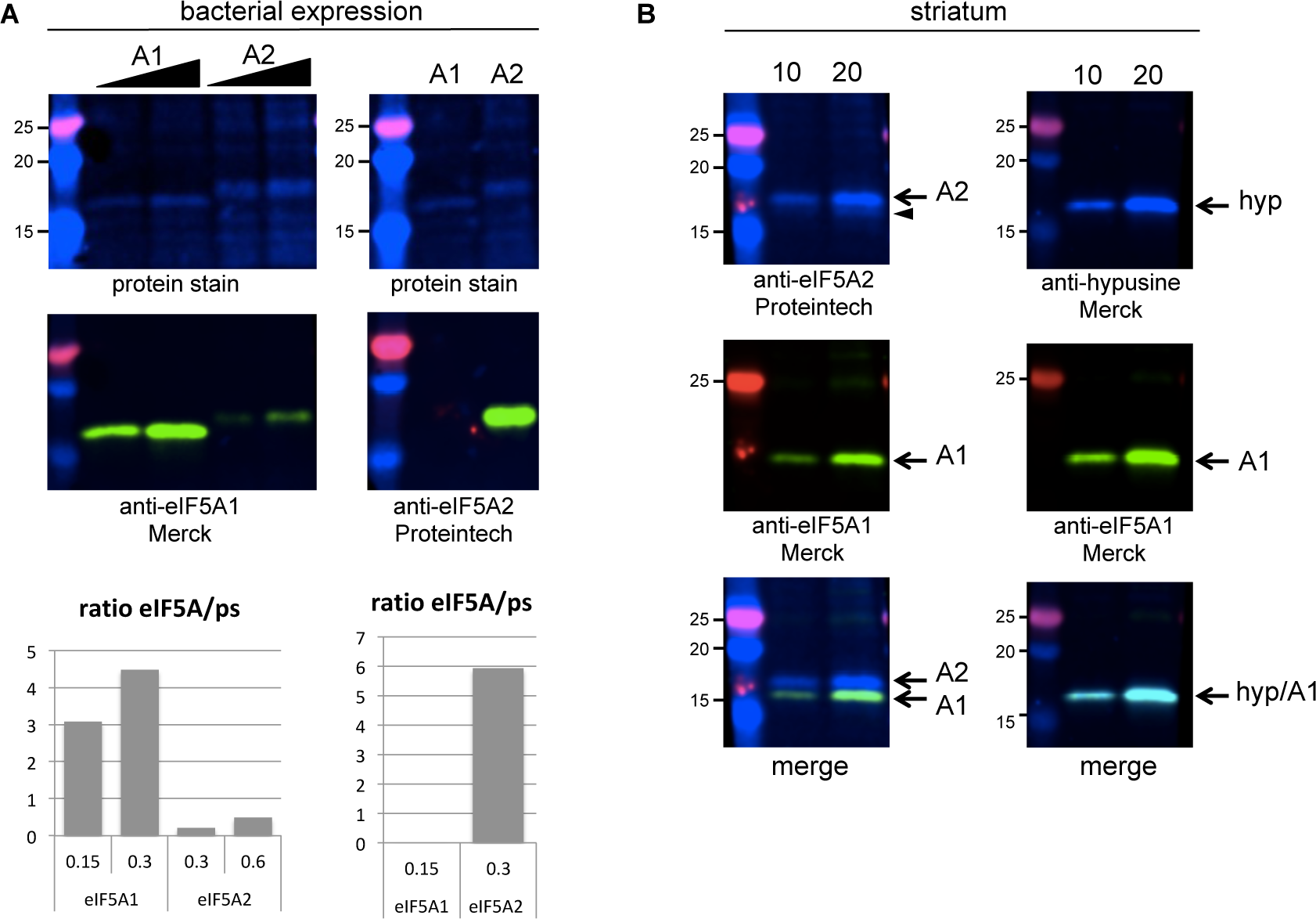
Specificity test of antibodies for eIF5A1 and eIF5A2. **(A)** The sequences coding for human eIF5A1 and eIF5A2 were amplified from the Human MTC Panel I (#636742, Clontech) and cloned in pRSET-B vector using NdeI and EcoRI sites and overexpressed in *Escherichia coli* C41 cells. Upper panels show total protein accumulation in C41 cells transformed with the corresponding plasmid either pRSET-eIF5A1 or pRSET-eIF5A2 (Revert™ 700 Total Protein Stain, 926-11021, LI-COR). Overexpressed eIF5A1 has a higher mobility than eIF5A2. Lower panels show the incubation of the above membranes with the corresponding antibodies. The mouse monoclonal antibody raised against eIF5A1 (SAB1402762, Merck) recognizes both proteins but preferentially recognizes eIF5A1. The quantification of the relative intensity normalized by the amount of the specific protein (protein stain of the corresponding band) showed that the antibody recognizes eIF5A1 approximately 10 times better than eIF5A2. Meanwhile, the rabbit polyclonal raised against eIF5A2 (17069-1-AP, Proteintech) efficiently recognizes eIF5A2 while very poorly recognizes eIF5A1 (the band is only visible if the membrane is overexposed). The quantification of the relatively intensity normalized to the amount of each corresponding protein shows that the antibody raised against eIF5A2 recognizes eIF5A2 200 times better than eIF5A1. See graphs on the bottom panels. **(B)** We evaluated the performance of these two antibodies in mouse striatal samples. Left-upper panel shows that the antibody against eIF5A2 recognizes a band of the expected size and a shadow band of higher mobility (arrow head, likely eIF5A1). The panel below shows that the antibody raised against eIF5A1 only recognizes a band of the expected size for eIF5A1 in the striatum. The merge of both channels (left-bottom panel) confirms that both proteins can be detected at the same time in the striatum and also confirms the higher mobility of eIF5A1 in striatal samples. These results also suggest, together with the quantifications made in (A) that the accumulation of eIF5A2 protein is lower than that of eIF5A1 in the striatum. On the right-upper panel we observed that the incubation of striatal samples with polyclonal anti-hypusine antibody (ABS1064-I, Merck) detects a single band and the co-incubation with mouse monoclonal antibody raised against eIF5A1 (SAB1402762, Merck) confirms that this band corresponds to eIF5A1 (merge bottom panel). Thus we cannot detect hypusinated eIF5A2 in the striatum, either because eIF5A2 does not have this modification or because the amount of hypusinated eIF5A2 does not reach the detection limits for the anti-hypusine antibody.

